# Privacy-Preserving Visualization of Brain Functional Connectivity

**DOI:** 10.1101/2024.10.11.617267

**Authors:** Ye Tao, Anand D. Sarwate, Sandeep Panta, Jessica Turner, Sergey Plis, Vince D. Calhoun

## Abstract

Data visualizations are an integral part of neuroimging research, supporting activities ranging from exploratory data analysis to the interpretation and communication of findings. While essential, visualizations can also reveal private information about individual participants. In this paper, we discuss how visualizations may inadvertently lead to privacy leakage and explore methods to mitigate such risks. Our work investigates ways to securely share visualizations that faithfully preserve the patterns supporting the derived insights from data analysis, rather than deriving conclusions from the visualizations themselves. We address the problem of privacy-preserving visualization under the framework of differential privacy, focusing on commonly used visualization methods for functional network connectivity. Several perturbation-based strategies are investigated for protecting correlation-related measures, with analyses of their privacy costs and the effects of pre- and post-processing. To achieve a better balance between privacy and visual utility, we propose workflows for connectogram and seed-based connectivity visualizations that preserve the qualitative structure of non-private results. Overall, this work illustrates how differential privacy can be effectively applied to neuroimaging visualization, highlighting its potential as a principled approach for safeguarding sensitive information.

## 1. Introduction

Data visualization (Kostelnick, 2008; Bhattacharjee et al., 2020) is an integral part of the research workflow, serving distinct purposes at different stages of analysis. In the early phase of exploratory data analysis, it enables researchers to examine data distributions and uncover underlying patterns. At later stages, visualizations facilitate the communication of analytical findings and aid in the interpretation of results. While privacy concerns are often discussed in the context of sharing raw data or summary statistics (Sweeney, 1997; Narayanan and Shmatikov, 2008), visualizations derived from these data can also expose sensitive information, particularly in domains involving human health and behavior, yet this aspect has received comparatively less attention than data leakage itself.

A range of privacy attacks have demonstrated that anonymized or aggregated outputs can still lead to re-identification or data leakage. Classic examples include linking de-identified health records with publicly available information Sweeney, 2002) and uncovering user identities from anonymized movie rating data (Narayanan and Shmatikov, 2008). Similar risks extend to visualizations, where adversaries may exploit graphical cues and/or external sources to infer sensitive information. A well-known real-world example is Strava’s release of an anonymized global activity heatmap, which unintentionally exposed secret U.S. military bases (Allen, 2020). Comparable risks arise in scientific visualizations. For example, scatterplots or boxplots commonly used in neuroimaging exploratory data analysis may reveal actual data values. Likewise, when a research team publishes average fMRI functional network connectivity (FNC) matrices for healthy and diseased groups, re-identification becomes possible: if an attacker knows the FNC of all subjects except one, the group membership of the remaining individual can be inferred by comparing the published visualizations (see Figure 1). Although this represents an extreme case, it illustrates how visualization can lead to privacy leakage, as auxiliary information about individuals may be accessible through various external sources or linkage attacks. This example also motivates the need for privacy-preserving visualization methods that remain robust even when an attacker possesses substantial background knowledge. By contrast, even when the attacker has only limited knowledge, published visualizations can still enable inference if privacy is not explicitly considered. Concretely, one can adapt membership inference tests (Homer et al., 2008; Dwork et al., 2015) to the visualization setting. Representing a published visualization as *v*_pub_ generated from a group, a target individual’s visualization as *v*_target_, and a reference sample as *v*_ref_, an adversary can compare similarity scores such as ⟨*v*_target_, *v*_pub_ ⟩ and ⟨*v*_ref_, *v*_pub_ ⟩, and determine whether the target belongs to the group based on their difference. Although such tests are abstract and rely on modeling assumptions, they demonstrate that releasing aggregate or visual summaries can still leak membership information in the absence of explicit privacy protection.

- We propose a series of novel methods for DP visualization in neuroimaging, including pre- and post-processing strategies to enhance visual quality.
- We systematically evaluate these methods across multiple real-world datasets, comparing the performance of different mechanisms and identifying their optimal composition strategies under varying privacy parameters.
- We develop end-to-end differentially private workflows for connectogram and SBC visualizations, illustrating group comparisons and significant connectivities.
- We assess the interpretability and utility of the proposed visualizations through both qualitative and quantitative analyses.

**Figure 1.**
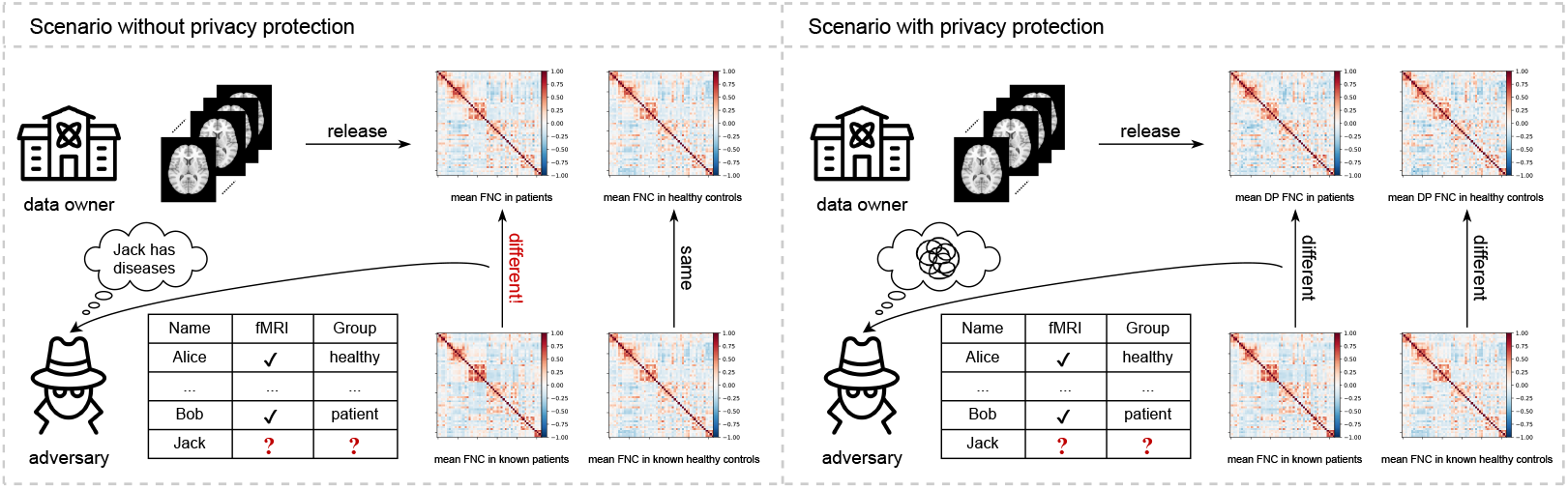
In a scenario without privacy protection, an adversary can infer information about unknown individuals. However, with differential privacy, the adversary cannot make such inferences.

Methods such as differential privacy (DP) (Dwork, 2006) offer protection against data leakage risks by introducing carefully calibrated randomness into the released outputs, thereby preventing accurate inference about any individual even when an adversary possesses substantial background knowledge. This protection often comes at the cost of a loss of accuracy and utility, which raises an important question: can privacy-preserving visualizations still capture the same patterns and convey the intended information? Previous study on differentially private visualizations, including boxplots, violin plots, and scatterplots (Tao and Sarwate, 2025; Panavas et al., 2023), has demonstrated that these methods can effectively protect individual information while preserving the essential patterns intended to be communicated (see Figure 2). Motivated by the growing need for privacy protection and the previous findings, this paper examines how privacy considerations can be incorporated into neuroimaging visualizations. Specifically, we explore the application of DP to commonly used visualizations, including FNC, connectogram, and seed-based connectivity (SBC) plots (Van Den Heuvel and Pol, 2010; Mohanty et al., 2020; Joel et al., 2011; Chopra et al., 2023). Our main contributions are summarized as follows.

**Figure 2.**
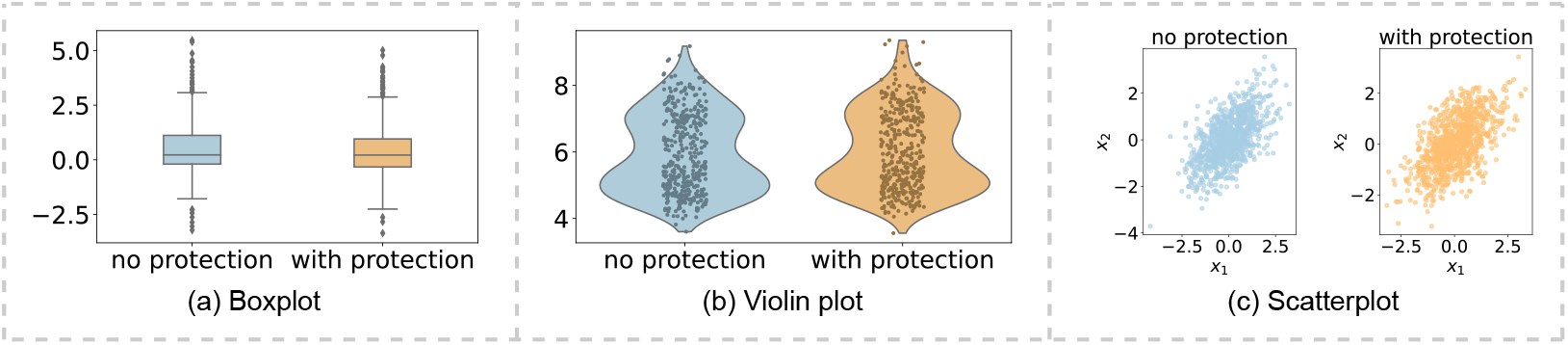
Comparison of visualizations with and without differential privacy protection, illustrating that differential privacy protects individual information while preserving the overall patterns and characteristics each plot is intended to communicate. Under differential privacy, the goal is to convey patterns rather than exact data values or make direct decisions; for example, boxplots show distribution and variability, violin plots reveal multimodal features, and scatter plots illustrate the increasing trend of one variable as the other grows.

## 2. Background

### 2.1. Differential Privacy

As data collection, storage, and analysis capabilities have improved significantly, privacy risks have increased, driving a growing focus on privacy research over the past two decades. Earlier syntactic approaches to privacy, such as de-identification and *k*-Anonymity (Garfinkel, 2015; Sweeney, 2002), are susceptible to linkage or membership inference attacks (Dwork et al., 2017; Merener, 2012; Ganta et al., 2008; Henriksen-Bulmer and Jeary, 2016; Fiore et al., 2020).

In contrast, differential privacy (Dwork et al., 2006; Dwork and Roth, 2014) has emerged as a state-of-the-art approach that provides a quantitative framework for privacy protection, addressing the vulnerabilities of earlier syntactic methods. It enables statistical analyses on datasets while ensuring that the privacy of individuals whose records are included is preserved. A differentially private algorithm or mechanism is a randomized function: the randomization is designed to make it difficult for an adversary observing the output to discern private information. The kind of information that DP protects is captured by the notion of neighboring databases. For two datasets *D* and *D*^′^ that are neighbors, denoted as *D* ∼ *D*^′^, the randomization should make it difficult for the adversary to distinguish, using a hypothesis test, whether the output was generated from *D* or *D*^′^ (Wasserman and Zhou, 2010; Kairouz et al., 2015). Most commonly, we say *D* ∼ *D*^′^ if they differ by the data of a single individual: this means that an adversary cannot reliably tell whether that individual was in the dataset or not. A full description of the key notations used in the paper is provided in Table 1. Some minor abuse of notation may occur, but it should be clear from the context.

**Table 1.** Key notations used in the paper.

| Notation | Description |
| --- | --- |
| $\epsilon$ | Privacy budget |
| $\delta$ | Probability of privacy violation or failure |
| $M$ | Randomized algorithm or mechanism |
| $D \sim D'$ | Neighboring datasets |
| $\Delta_1, \Delta_2$ | Global $\ell_1$-sensitivity and $\ell_2$-sensitivity of $f(D)$ |
| $\Delta_u$ | Sensitivity of the utility function $u$ |
| $v$ | Brain voxel |
| $r$ | A single ROI or $r$ brain regions, as defined by context |
| $t, T$ | Timepoint and the total number of timepoints, respectively |
| $S(\cdot, t)$ | BOLD value for a voxel or an ROI at time $t$ |
| $X_{ij}$ | Correlation coefficient between the timeseries of $r_i$ and $r_j$ |
| $X^{(n)} \in \mathbb{R}^{r \times r}$ | FNC matrix for subject $n$ |
| $\bar{X}, \tilde{X}$ | Average FNC and perturbed FNC |

For an algorithm to be differentially private, it must guarantee that the adversary’s test is unreliable for all possible *D* ∼ *D*^′^. This unreliability is measured in DP using two parameters, *ϵ* and *δ* (see Definition 1). The parameter *ϵ* is the privacy budget that quantifies the maximum distance between the output distributions of algorithm *M* when it runs on any two neighboring datasets. A smaller *ϵ* will give stronger privacy but less accurate responses. The parameter *δ* represents the probability of information leakage. Typically, we are interested in values of *δ* that are far less than the inverse of the database size, *δ* ≪ 1*/N* . The definition of DP ensures that for all neighboring datasets *D* and *D*^′^, the absolute value of the privacy loss (Dwork and Roth, 2014) is bounded by *ϵ* with a probability of at least 1 — *δ*.

#### Definition 1

((*ϵ, δ*)**-DP (Dwork and Roth, 2014))**. *Let ϵ, δ* ≥ 0, *a randomized algorithm M* : 𝒟 → ℋ *is* (*ϵ, δ*)*-differentially private if for all* 𝒴 ⊆ ℋ *and for all neighboring datasets D, D*^′^ ≈ 𝒟,

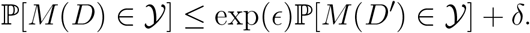

To design a (*ϵ, δ*)-DP algorithm that approximates a desired function *f* : 𝒟 → ℝ^d^ (in our case, a data visualization), a common approach is to add random noise of appropriate magnitude to the output of *f* (*D*). The amount of noise required depends not only on *ϵ* and *δ* but also on the sensitivity of the function, which quantifies the maximum change in the output when a single data point in the dataset is modified. The global *l*_1_ and *l*_2_ sensitivity of the function *f* are given by

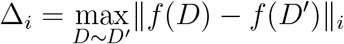

for *i* = 1 or 2, where ∥ *·* ∥_i_ denotes the *l*_i_-norm. “Pure” (*ϵ*, 0)-DP can be achieved by applying the Laplace mechanism (Dwork and Roth, 2014), defined as *M*_L_(*D*) = *f* (*D*) + *Z*, where *Z* follows the Laplace distribution *Z* ∼ Laplace(0, Δ_1_*/ϵ · I*). The identity matrix *I* ∈ ℝ^d^ is included when *f* (*D*) is high-dimensional. Alternative noise distributions have also been proposed (Geng et al., 2015; Alghamdi et al., 2022b). For any *ϵ, δ* ≈ (0, 1), the classical Gaussian mechanism *M*_G_(*D*) guarantees “approximate” (*ϵ, δ*)-DP when *M*_G_(*D*) = *f* (*D*)+*Z*, where *Z* ∼ 𝒩 (0, *σ*^2^*·I*) and 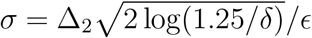. However, this choice of *σ* can be overly conservative in the high privacy regime (*ϵ* → 0) and does not apply in the low privacy regime (*ϵ* → ∞). To address this, Balle and Wang (2018) proposed an “analytic” formula for the variance.

In the following sections and experiments, we use the analytic Gaussian mechanism rather than the classical Gaussian mechanism when considering noise from the Gaussian distribution.

Additive noise mechanisms are well-suited for privatizing real-valued functions. In other scenarios, algorithms may instead randomly select an output from a set of choices. In such cases, the exponential mechanism (McSherry and Talwar, 2007) enables (*ϵ*, 0)-DP randomized selection from a discrete set. Sometimes, numerical outputs can be mapped to a discrete set. For example, in visualization, certain numerical values are mapped to the same color. Given a range ℋ, the exponential mechanism is defined with respect to a utility function *u* : 𝒟 × ℋ →ℝ, which assigns a utility score to each input-output pair. The exponential mechanism *M*_E_(*D*) selects and outputs an element *H* ≈ ℋ with a probability proportional to 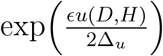, where 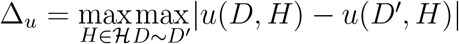.

Mechanisms such as additive noise or randomized selection protect privacy in a single analysis. When applied multiple times, the accumulated privacy loss can be formally quantified using the composition property of DP. There are several types of composition rules: basic, advanced, exact, and Rényi differential privacy (RDP)-based compositions (Dwork et al., 2010; Dwork and Roth, 2014; Kairouz et al., 2015; Murtagh and Vadhan, 2015; Abadi et al., 2016; Mironov, 2017; Balle et al., 2020; Dong et al., 2022; Sommer et al., 2018; Alghamdi et al., 2022b,a; Koskela and Honkela, 2020; Gopi et al., 2021; Koskela et al., 2020; Ghazi et al., 2022; Doroshenko et al., 2022). A full discussion of composition rules is beyond the scope of this paper, but for the purposes of this work, advances in composition analyses can only improve the guarantees that we make. Another important property of DP is post-processing immunity: any function applied to the output of a differentially private mechanism cannot weaken its privacy guarantees. This property ensures that subsequent analyses or transformations remain safe without additional noise.

### 2.2. Privacy-preserving Visualizations

#### Anonymization and perturbation-based approaches

Early studies, such as Wang et al. (2018) and Chou et al. (2017), adopted an “anonymization” model to guarantee privacy. Such syntactic models, however, can still leak information through composition attacks (Narayanan and Shmatikov, 2008; Dwork and Roth, 2014; Ganta et al., 2008). Avraam et al. (2021) later proposed probabilistic anonymization, which introduces random noise to individual values for creating privacy-preserving visualizations. While conceptually related to DP, these methods do not quantitatively measure privacy risk.

#### Algorithmic approaches to differentially private visualization

Differential privacy provides formal guarantees even in the presence of side information and has become the gold standard for privacy protection. Zhang et al. (2016) were among the first to systematically examine the challenges of privacy-preserving data visualization. Using two-dimensional location data as an example, they constructed grids over the data domain and applied the Laplace mechanism to compute noisy counts for each grid cell. Their work analyzed visual artifacts introduced by noise and proposed ways to communicate uncertainty in private visual outputs. Building on this work, Zhang et al. (2020) extended the Laplace mechanism to various visualization types, including bar charts, and designed strategies to adjust the amount of noise for improving visualization utility. This approach mitigates distortion while maintaining privacy guarantees. In a related study, Lee (2017) qualitatively examined the effects of Laplacian noise on various visualization types, such as bar graphs, pie charts, heatmaps, linear plots, and scatterplots. Using smart grid electricity consumption time-series data, the study explored how DP constraints affect visual cues and identified pre- and post-processing methods for restoring them.

#### Evaluation of visual perception and utility under DP

Panavas et al. (2023) focused on evaluating the perceptual utility of differentially private scatterplots. Their comparative study showed that factors such as algorithm choice (e.g., AHP, DAWA), privacy level, bin number, data distribution, and user task all influence the visual outcome. To address the lack of guidance in balancing these factors, they had experts evaluate 1200 DP scatterplots to assess their effectiveness in preserving aggregate patterns. The results established a ground truth for visual utility and identified multi-scale structural similarity (MS-SSIM) as an effective objective metric for optimizing parameter selection in DP scatterplots. Liu et al. (2019) examined the common belief that aggregating individual data into composite representations such as heatmaps protects privacy. However, their analysis of noise-free heatmaps using eye-tracking data showed that they do not guarantee privacy. To address this, they proposed two noise mechanisms that ensure privacy and analyzed their privacy-utility tradeoff. Their results indicated that the Gaussian noise mechanism, compared to the Laplace mechanism, provides a more effective solution for preserving privacy in heatmaps. These findings have significant implications for the development of differentially private mechanisms in the eye-tracking domain.

#### Extensions to specialized visualization types

More recent work, motivated by Liu et al. (2019) and Steil et al. (2019), has looked at specific visualizations such as heatmaps and boxplots. Ghazi et al. (2023) presented an efficient DP algorithm for heatmaps. The core of their algorithm was a DP procedure that took a set of distributions as input and produced an output close to the average of the inputs in terms of earth mover’s distance (EMD). EMD better captures perceptual similarity compared to other measures such as *l*_1_, *l*_2_, or Kullback-Leibler (KL) divergence. They also proved theoretical bounds on the error of their algorithm under a sparsity assumption, demonstrating that these bounds were near-optimal. Ramsay and Diaz-Rodriguez (2024) introduced a differentially private boxplot and evaluated its effectiveness in displaying location, scale, skewness, and tails of a given empirical distribution. They theoretically demonstrated that the location and scale of the boxplot are estimated with optimal sample complexity, and skewness and tails are estimated consistently. This is not always the case for a box-plot naively constructed from a single existing differentially private quantile algorithm.

#### System-level frameworks for DP visualization

Interactive systems for private data visualization have been proposed by Nanayakkara et al. (2022) and Budiu et al. (2022), focusing on the relationship between privacy budget and accuracy to support *data privacy management* rather than visualization itself. Zhou et al. (2022) introduced the PriVis model, a Bayesian network-based differential privacy framework, to generate private scatter, line, and bar charts while preserving user-preferred patterns, addressing a limitation of prior work, which often targeted specific visualization types or algorithms without systematically considering the importance of data patterns in the visual output. They also developed the DPVisCreator system to assist data custodians in implementing this approach and evaluated its effectiveness through quantitative analysis of pattern utility, case studies, and semi-structured expert interviews.

#### Limitations of prior work and motivation for our study

Despite these advances, prior work mainly focused on generic visualization types rather than domain-specific cases such as brain functional connectivity, which may require different techniques. Most studies do not systematically compare DP mechanisms to minimize required noise, and pre- and/or post-processing methods often lack structured workflows, limiting their applicability to neuroimaging. Evaluations also rely heavily on qualitative assessment without exploring potential quantitative approaches to measure visual utility. In contrast, our work focuses on functional connectivity in neuroimaging and provides a systematic evaluation of DP mechanisms, composition, and pre- and/or post-processing methods.

### 2.3. Functional Connectivity in Neuroimaging

In this work, we focus on neuroimaging data, particularly functional magnetic resonance imaging (fMRI), due to its sensitivity, high dimensionality, and collaborative use across institutions. fMRI data can reveal aspects of an individual’s cognitive state, making privacy protection critical. Differential privacy offers a principled framework for secure data sharing in such settings. Furthermore, the variety of visualization modalities employed in fMRI studies enables a comprehensive examination of privacy-preserving visualization techniques, with insights that can extend to other neuroimaging modalities and scientific domains.

Resting-state functional MRI (rs-fMRI) captures the blood oxygenation level dependent (BOLD) signal while subjects remain in the scanner without performing specific tasks. A series of MRI images is acquired over time, with each image recording BOLD signal intensity at each voxel. We denote the BOLD value for voxel *v* at time *t* as *S*(*v, t*). Typical MRI images contain roughly 100, 000 voxels, which can be grouped into regions of interest (ROIs) using various parcellation schemes (Tzourio-Mazoyer et al., 2002; et al., 2012; Gordon et al., 2016; Du et al., 2020). ROI timeseries are obtained by averaging voxel-level signals within each ROI: 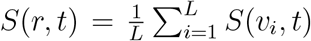, where *r* is the ROI and *L* is the number of voxels in that ROI. Analyses at the voxel and ROI levels provide complementary insights into brain functional connectivity, motivating diverse visualization approaches to effectively present these results.

FNC quantifies correlations between activity in different brain regions ((Mohanty et al., 2020). Various methods exist to compute these correlations (Marrelec et al., 2006; Hlinka et al., 2011); in this work, we typically use the Pearson correlation coefficient, which ranges from —1 to 1 and measures the linear relationship between ROI-based timeseries. For any two ROIs, the correlation is given by

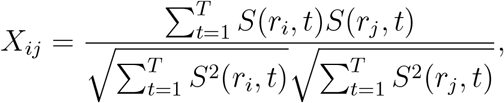

where *T* denotes the number of timepoints and *S*(*·, t*) is assumed to be centered at 0. The FNC for a subject *n* across *r* brain regions is represented by an *r* × *r* adjacency matrix *X*^(n)^, with each entry corresponding to the correlation between a pair of regions. We focus on visualizing the average FNC across a group of *N* subjects (see Section 3.1), defined as

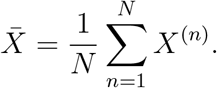

For visualization, the elements of 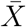 are normalized to the range [0, 1], quantized to a set of color values 𝒞, and mapped to RGB triples (*R*_ij_, *G*_ij_, *B*_ij_). In addition to visualizing the average FNC matrix, a connectogram can be used to represent FNC results graphically, with nodes as brain regions and edges indicating connection strength (see Section 3.2).

While FNC captures large-scale network interactions using ROI-based timeseries, SBC provides detailed information about the connectivity of a specific seed region with the rest of the brain. SBC involves selecting a seed voxel or ROI and computing its correlation with the timeseries of all other brain voxels. The resulting connectivity patterns are typically visualized as SBC spatial maps (Joel et al., 2011) (see Section 4).

### 2.4. Challenges in DP Functional Connectivity Visualization

There are (at least) three potential issues to consider when using differential privacy for visualization: the mismatch between the specific computations needed in the visualization and existing (generic) DP methods, the issues of multiple data accesses coming from applications in exploratory data analysis, and the challenge of defining the utility of data visualizations (Bhattacharjee et al., 2020; Zhang et al., 2016; Zhou et al., 2022). When thinking about visualization, the notion of “utility” can differ from standard quantitative metrics. In some scenarios, a private visualization has high utility if an expert would draw the same qualitative conclusions as from the non-private (unprotected) visualization.

To address these challenges, we take the following approaches. For the computation mismatch between specific visualization tasks and existing generic DP methods, we adapt DP mechanisms specifically for FNC to provide a private estimate of the correlation matrix underlying the visualization. In the case of the connectogram, we introduce a DP workflow designed for group comparisons, and for SBC visualization, we propose a DP procedure to systematically identify and present statistically significant results. To manage multiple data accesses, we employ (adaptive) composition techniques (Kairouz et al., 2015; Dong et al., 2020, 2022; Mironov, 2017) to quantify and control cumulative privacy risk. Finally, to address the challenge of utility, we provide both qualitative assessments and quantitative measurements.

In addressing these issues, we outline a systematic process for designing privacy-preserving visualizations across different visualization types and application domains, emphasizing both visualization quality and data privacy. Although our focus is on neuroimaging, the *process* we outline can be extended to other domains. Visualization typically involves computing or aggregating statistics from raw data through multiple steps. The first step is to identify the statistics or aggregated quantities needed for visualization. Next, suitable pre-processing strategies are applied to reduce the noise required for privacy protection. Different visualization types may benefit from different pre-processing techniques. We then design the differentially private mechanism, which may involve selecting appropriate noise distributions, applying composition methods, or constructing privacy-preserving workflows. Finally, post-processing methods are used to mitigate the effects of perturbation. Following this process, we demonstrate how privacy-preserving visualizations are constructed in neuroimaging (see Sections 3 and 4).

## 3. FNC Visualization

### 3.1. FNC Matrix Visualization

In this section, we describe the privacy-preserving visualization process for the average FNC matrix, summarized in Table 2, together with pre- and post-processing techniques that enhance visual utility.

**Table 2.** A summary of the mechanisms explored in this paper is provided. Although other noise distributions may perform better for specific tasks, we focus on Laplace and Gaussian mechanisms, which are the most widely studied, for clarity of exposition.

| Algorithm | Name | Strategy | Mechanism | Composition |
| --- | --- | --- | --- | --- |
| matrix-level | Pure | add perturbation | Laplace | — |
|  | Approx | add perturbation | Gaussian | — |
|  | Exp | random mapping | Exponential | — |
| entry-level | Pure<sub>k</sub> | add perturbation | Laplace | exact |
|  | Pure<sub>r</sub> | add perturbation | Laplace | RDP-based |
|  | Approx<sub>k</sub> | add perturbation | Gaussian | exact |
|  | Approx<sub>r</sub> | add perturbation | Gaussian | RDP-based |
|  | Exp<sub>k</sub> | random mapping | Exponential | exact |

#### 3.1.1. Matrix-level Differentially Private Algorithms

In FNC matrix visualization, the true outputs are the RGB triples corresponding to each pairwise correlation in the average matrix. To guarantee DP, we adopt the following strategy: first compute the average FNC matrix under DP, then apply standard quantization and color mapping. This strategy falls under *output perturbation*, where noise is added directly to the average FNC matrix:

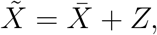

where the noise distribution *Z* is determined by the privacy parameters. After perturbation, the noisy outputs 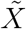 are truncated to the valid range [—1, 1]. We consider two mechanisms:

- Pure: (*ϵ*, 0)-DP Laplace mechanism. Let *Z* ∼ Laplace(0, Δ_1_*/ϵ · I*), where Δ_1_ = *r*(*r* − 1)*/N* is the *l*_1_ sensitivity of 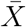.
- Approx: (*ϵ, δ*)-DP analytic Gaussian mechanism. Let *Z* ∼ 𝒩 (0, *σ*^2^ *· I*), where *σ* is the minimum standard deviation that satisfies the condition in (Balle and Wang, 2018, Theorem 8), based on the *l*_2_ sensitivity 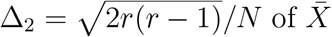.

#### 3.1.2. Entry-level Differentially Private Algorithms

For the matrix-level methods, we first calculate the overall sensitivity of the FNC matrix and then generate the corresponding perturbed matrix. More efficient approaches may be obtained by applying composition methods at the entry level. Given total privacy parameters (*ϵ, δ*) and sequence size *k* = *r*(*r* — 1)*/*2, we consider the following variants:

- Pure_k_: We compute the per-entry privacy parameters (*ϵ*^′^, 0) using exact composition and apply the Laplace mechanism to each entry with sensitivity Δ = 2*/N* . Hence, 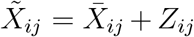, where *Z*_ij_ ∼ Laplace(0, Δ*/ϵ*^′^). For one-dimensional outputs, the global *l*_1_ and *l*_2_ sensitivities are identical, so we use Δ to denote the sensitivity without distinction between norms.
- Pure_r_: Given the overall privacy guarantee, we compute the scale parameter *b* of the Laplace distribution using RDP-based composition. Then, 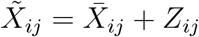, where *Z*_ij_ ∼ Laplace(0, *b*).
- Approx_k_: We compute the per-entry privacy parameters (*ϵ*^′^, *δ*^′^) using exact composition and apply the analytic Gaussian mechanism with sensitivity Δ = 2*/N* . Thus, 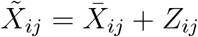, where the variance of *Z*_ij_ satisfies the inequality in (Balle and Wang, 2018, Theorem 8).
- Approx_r_: We compute the variance *σ*^2^ of the Gaussian distribution using RDP-based composition under the given privacy parameters. Then, 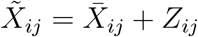, where *Z*_ij_ ∼ 𝒩 (0, *σ*^2^).

Visualization often serves as a preparatory step in research or exploratory data analysis, which may require accessing raw data multiple times and generating visualizations from different intermediate statistics. In such cases, (adaptive) composition plays a critical role. The matrix-level Laplace mechanism provides a strong “pure” privacy guarantee (*δ* = 0), but by allowing a slightly weaker guarantee (*δ >* 0), we can enhance the privacy budget through composition methods, reducing the level of noise required. For the Gaussian mechanism, we compare the performance of matrix-level and entry-level analytic Gaussian mechanisms.

We also explored the Exponential mechanism (McSherry and Talwar, 2007), which performs a randomized mapping from 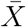 to the set of all RGB images ℋ, either as a whole (denoted Exp) or entry-by-entry (denoted Exp_k_). The Exp method provides (*ϵ*, 0)-DP by sampling *H* ≈ 𝒩 according to the utility function 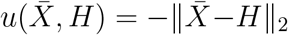, but this is computationally infeasible due to the size of 𝒩. The entry-level approach Exp_k_ applies an (*ϵ*^′^, 0) mechanism with exact composition to map each 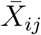 to a color *C* ≈ 𝒞 using the utility function 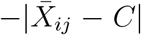 with sensitivity 2*/N* . However, this approach yielded substantially poorer visual quality compared to output perturbation–based methods.

##### Algorithm 1

Method for choosing the clipping bound *b*

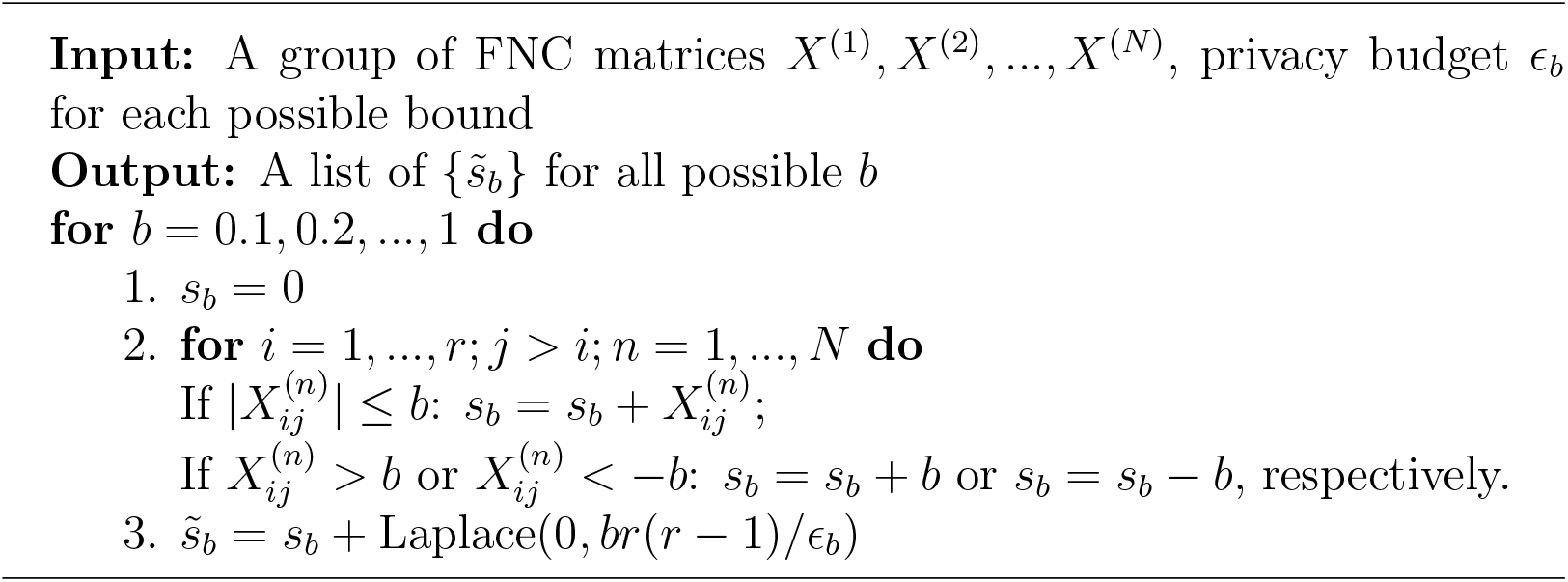

#### 3.1.3. Pre-processing Approaches

Given privacy parameters *ϵ* and *δ*, some pre-processing approaches can help reduce the required noise. One effective approach is range reduction. Prior information of FNC suggest that extremely high correlations (e.g., 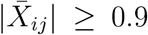,*i* /= *j*) are rare. Thus, the range of 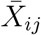 can be constrained to [—*b, b*], where 0 *< b <* 1. Under this clipping, the *l*_1_ and *l*_2_ sensitivities of 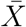 are Δ_1_ = *br*(*r* − 1)*/N* and 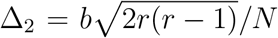, respectively, and each entry 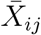 has sensitivity Δ = 2*b/N* .

In this work, we employ a clipping method (Near and Abuah, 2021) to estimate the bounds of the FNC range (see Algorithm). The method establishes upper and lower bounds for 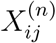 by gradually increasing the bound *b* from 0 to 1. If 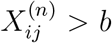 or 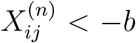, the value is clipped to *b* or —*b*, respectively. We then calculate the sum of all entries for each possible bound and apply the Laplace mechanism to ensure the privacy of the total sum. Finally, we select the bound that yields a stable perturbed sum (see Figure 3).

**Figure 3.**
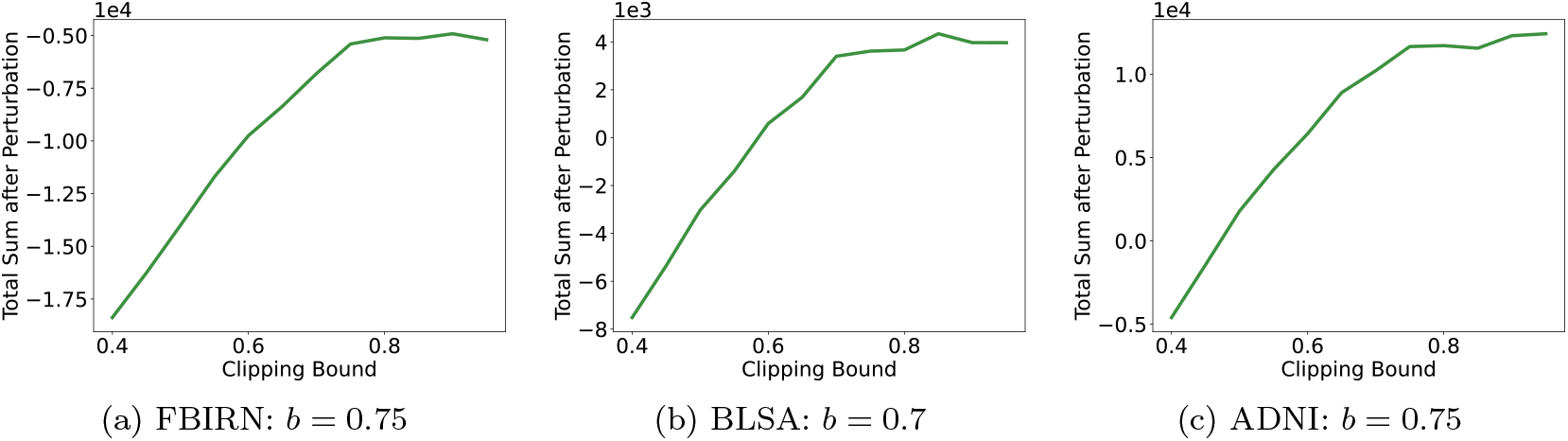
Clipping method for choosing *b* on three datasets. The total privacy budget allocated for bound estimation is 0.5. The search for *b* begins at 0.4 rather than 0, assuming that the bound of FNC values exceeds 0.4.

**Figure 4.**
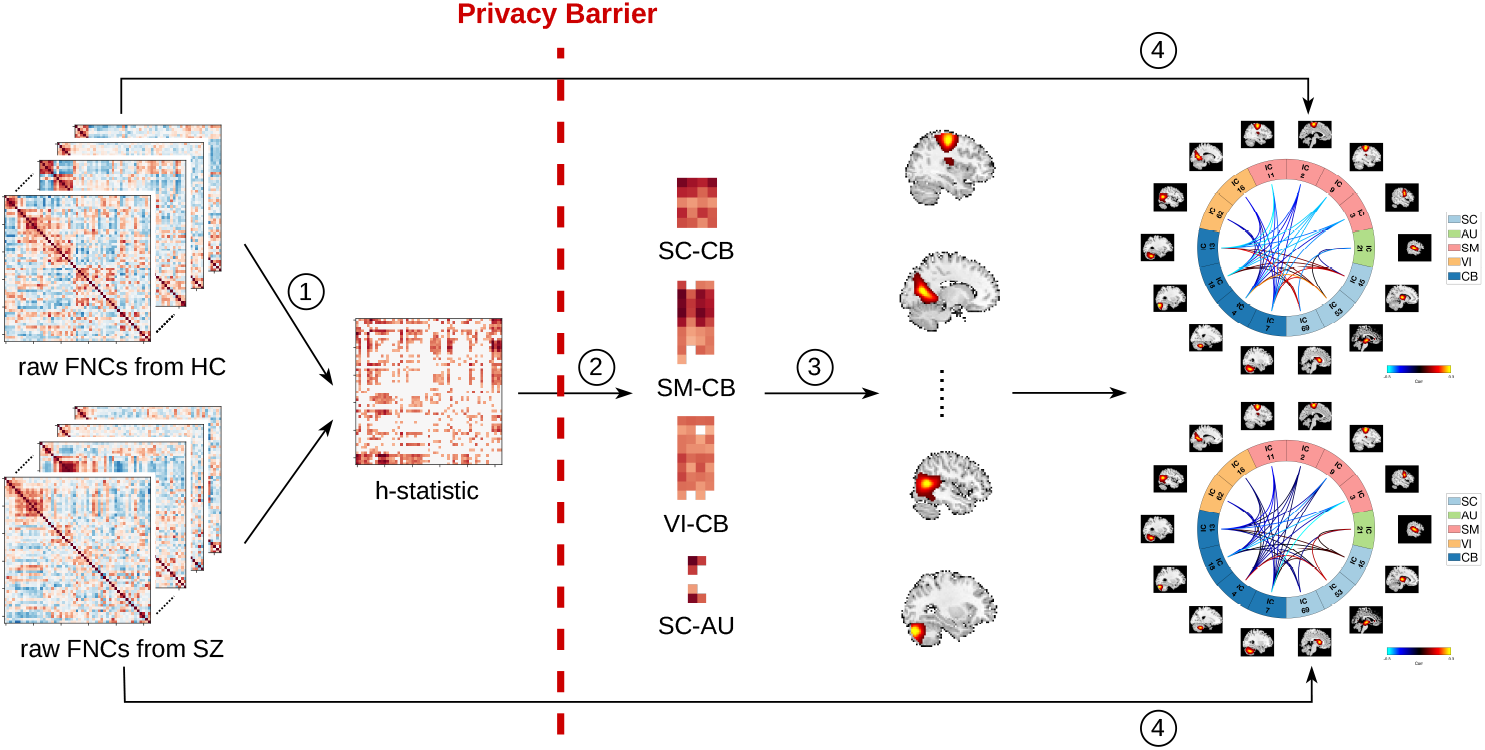
Privacy-preserving workflow for connectogram visualization. The privacy barrier refers to the stage where sensitive data (e.g., FNC values, *h*-statistics) are perturbed using differential privacy techniques. Data after the privacy barrier is considered privacypreserving, and it is impossible to infer the true data from this released information.

Alternative techniques can be used to estimate the bounds, such as leveraging prior domain knowledge, using public data, or applying different DP methods. For example, public datasets can help determine upper and lower bounds. Another approach is to estimate differentially private quantiles and use them as bounds (McSherry and Talwar, 2007; Smith, 2011). Liu et al. (2019) propose an algorithm that optimizes the expected mean squared error between perturbed aggregated data and its original counterpart to estimate bounds. Regardless of the method used, there is a tradeoff between information loss and the noise required for differential privacy. When the upper and lower bounds are set closer to each other, less noise is needed to guarantee differential privacy due to a lower sensitivity. However, using smaller clipping bounds results in the removal of a significant amount of information.

#### 3.1.4. Post-processing Approaches

Differential privacy has a post-processing invariance property, meaning that any data-independent operations on differentially private outputs do not incur additional privacy loss. Hence, post-processing can be used to “clean up” images without compromising privacy. This can help mitigate the effects of perturbation on FNC or any other visualizations and potentially improve visual utility.

For FNC visualization, we explore two post-processing approaches: Singular Value Decomposition (SVD) and Haar wavelet transform (Mulcahy, 1997). The SVD of the differentially private output is 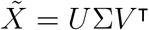. We select the first *s* components after examining the magnitudes of the singular values. If the added noise does not destroy the visualization pattern, we are able to observe that the first *s* singular values are significantly larger than the remaining ones. For visualization, we use 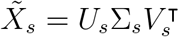 after post-processing. In the Haar wavelet transform, we apply a transformation to obtain the matrix 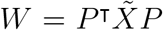, where *P* is the invertible Haar transformation matrix. In many cases, the wavelet-transformed matrices are sparser than their original counterparts. We then threshold the coefficients in matrix *W* to generate a new coefficient matrix, denoted as *W*_w_. Finally, we perform the inverse wavelet transform on *W*_w_ to obtain 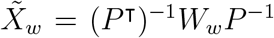, which represents the processed data used for visualization.

### 3.2. Connetogram Visualization

#### 3.2.1. Non-private Workflow for Group Comparison

Connectogram visualizations effectively represent group differences in brain regions and functional connectivity, such as comparisons between healthy controls (HC) and schizophrenia (SZ) patients. The standard workflow for generating a connectogram first transforms each element of the FNC matrix into Fisher’s *Z*-scores. The statistical significance of connectivity strength differences between HC and SZ groups is then assessed using a two-tailed two-sample *t*-test (*p <* 0.05 with Bonferroni correction) (Du et al., 2020).

#### 3.2.2. Private Workflow for Group Comparison

We propose a novel differentially private workflow for connectogram visualization. A straightforward approach might be to add perturbation directly to the *p*-values or *t*-statistics. However, this is challenging because the sensitivity of the *t*-statistic is complex to compute, and added noise can substantially alter the *p*-values. Our experiments indicate that, under differential privacy, nonparametric tests (Couch et al., 2019) are more effective for constructing connectograms. Consequently, we adopt a private workflow based on the nonparametric absolute Kruskal-Wallis (AKW) test.

Consider a database of size *N* partitioned into *g* groups. Let *N*_i_ denote the size of group *i*, and *r*_ij_ the rank of the *j*-th element in group *i*. Define the mean rank of group *i* as 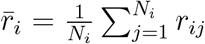, and the overall mean rank as 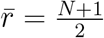. The absolute Kruskal-Wallis *h*-statistic is given by

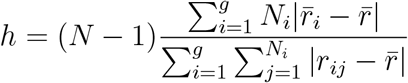

and has a sensitivity of 8 (Couch et al., 2019, Algorithm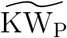).

Our workflow (see Figure) proceeds as follows: (1) Compute the AKW test on FNC matrices from two groups to obtain an *h*-statistic matrix. (2) For each *h*-statistic submatrix, which is partitioned by brain domains (Du et al., 2020, Table 2), calculate the average *h*-statistic. Apply the Report Noisy Max (RNM) algorithm (Dwork, 2006) to select the top *p*_d_ submatrices in a differentially private manner. The RNM algorithm is a differential privacy technique for private selection. Instead of selecting the top-ranked items based on their true values, RNM adds calibrated noise to each value and selects the top-ranked items based on the noisy values. The number of selected items is determined by the privacy budget allocated at this stage. (3) Average the *h*-statistic for each brain region within the selected brain domains and apply RNM again to select the top *p*_r_ brain regions privately. (4) Finally, compute perturbed connectivities from the average FNC of each group. With privacy-preserving nodes and edges, the connectogram for group comparisons can be visualized.

## 4. SBC Visualization

### 4.1. Non-private Workflow for Significant Connectivity

SBC characterizes the connectivity pattern between a predefined seed and the entire brain. This involves computing Fisher-transformed bivariate correlation coefficients between the seed BOLD timeseries and the timeseries of each individual voxel (Joel et al., 2011). The correlation between the seed and any voxel *v* is defined as

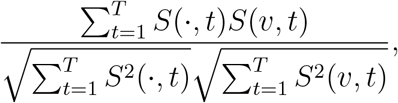

where *S*(*·, t*) represents the BOLD timeseries from the seed (either a single voxel or an ROI) at time *t*, and *T* is the number of timepoints in the experiment. The seed timeseries *S*(*·, t*) is assumed to be mean-centered. The correlation coefficient is then converted to a *z*-score using the Fisher *Z*-transform. A one-sample *t*-test is subsequently performed to determine whether the mean connectivity across subjects significantly differs from zero, indicating significant positive or negative connectivity.

### 4.2. Private Workflow for Significant Connectivity

The privacy-preserving SBC visualization process is quite similar to connectogram visualization and also involves two main components (see Figure 5): selecting significant connectivities in a privacy-preserving manner and displaying *z*-scores with differential privacy. To identify statistically significant connectivities, we employ the Wilcoxon signed-rank (WSR) test, a nonparametric alternative to the one-sample *t*-test, as nonparametric tests are often more effective than their parametric counterparts. For a given database, let *s*_i_ denote the sign of the data *x*_i_, and let *r*_i_ represent the rank of |*x*_i_|, determined by the magnitude of the data. The WSR test statistic is then calculated as *w* = ∑*s*_i_*r*_i_. Connectivities are selected at a significance level of *p <* 0.05 after applying Bonferroni correction (Couch et al., 2019, Algorithm 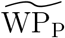).

**Figure 5.**
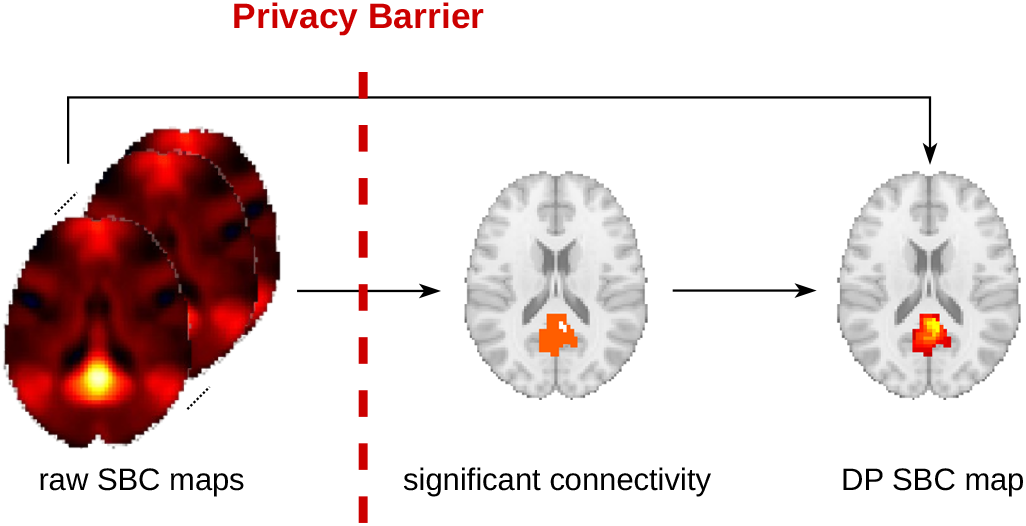
Privacy-preserving workflow for SBC map visualization.

To protect the *z*-scores, we use the analytic Gaussian mechanism to generate noise, which is added to the correlations before applying the Fisher *Z*-transform. Although narrowing the range of correlations could reduce the required noise, this approach is not suitable here because high correlations are expected (see Figure 6a). The privacy-protected correlation map exhibits noticeable salt-and-pepper noise (see Figure 6b), which is absent in FNC visualization. One likely reason is that connections within the same brain network tend to be strong, resulting in relatively stable and similar correlation values. To mitigate this noise, we apply a median filter (Erkan et al., 2018) as a post-processing method (see Figure 6c). After identifying significant connectivities and their corresponding values, we construct the privacy-preserving SBC map.

**Figure 6.**
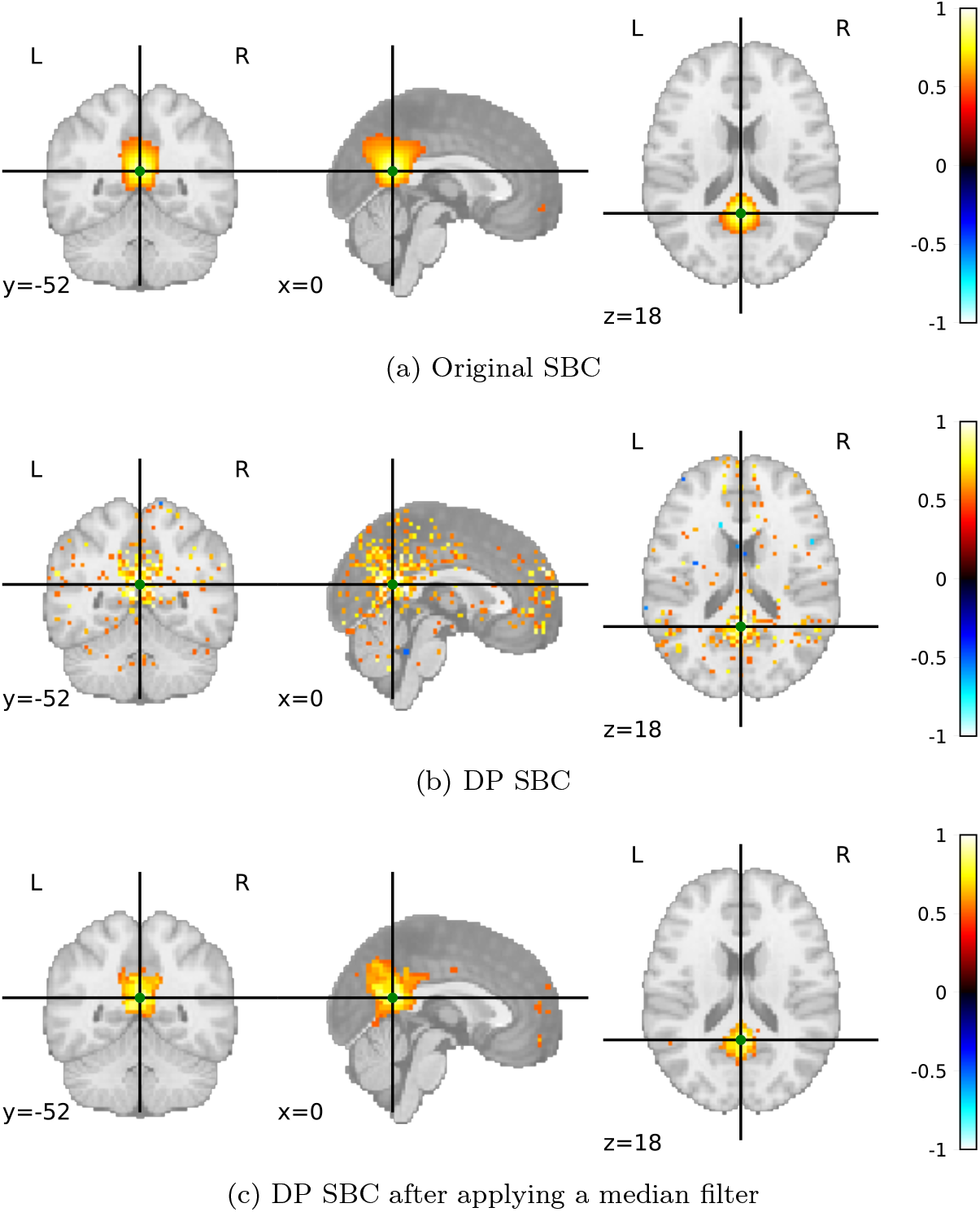
SBC maps on the NITRC dataset using the Posterior Cingulate Cortex (PCC) as the seed (indicated by the green dot), with parameters *N* = 1082, *ϵ* = 2.5, and *δ* = *N*^*—*3*/*2^. Correlations with absolute values greater than 0.5 are displayed.

## 5. Experiments and Results

We aim to investigate the following research questions: (1) Which algorithm most effectively preserves functional connectivity patterns under the same level of privacy? (2) Can pre- and post-processing methods enhance visualization quality? (3) Does the differentially private connectogram workflow identify significant brain regions and connections as effectively as the non-private workflow? (4) Is the differentially private workflow for SBC visualization as effective as its non-private counterpart in identifying significant connectivities? (5) How do numerical utility measures, such as mean squared error (MSE), relate to visualization quality?

We conducted privacy-preserving FNC visualization experiments using datasets from the Function Biomedical Informatics Research Network (FBIRN), the Baltimore Longitudinal Study of Aging (BLSA), and the Alzheimer’s Disease Neuroimaging Initiative (ADNI) (Du et al., 2020; Sendi et al., 2021; Ferrucci, 2008). Our findings were consistent across all datasets. For connectogram visualizations, we used the FBIRN dataset to compare HC and SZ groups. For SBC visualizations, we employed [NITRC](https://www.nitrc.org/plugins/mwiki/index.php/neurobureau:AthenaPipeline#Whole_Brain_Data) the dataset.

### 5.1. Methods Comparison

In general, we found that Gaussian mechanisms preserved visual utility better than Laplace mechanisms, which in turn outperformed Exponential mechanisms (see Figure 7, Figure A.3, and Figure A.4). Overall, Approx-Clip-SVD achieved the best performance.

**Figure 7.**
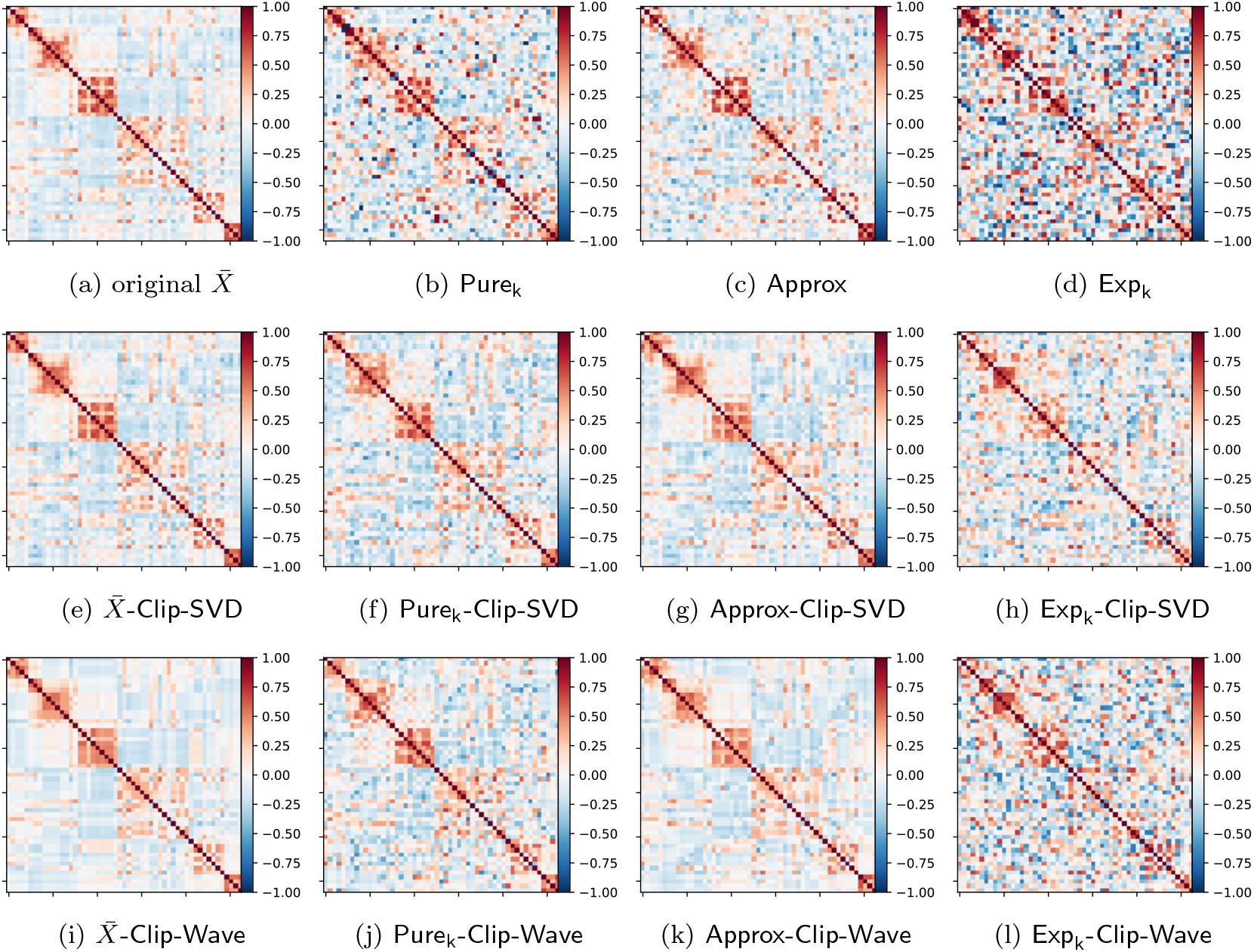
Visualization of the FBIRN dataset with *N* = 1056, clipping bound *b* = 0.75, number of components used in SVD set to 10, threshold used in the wavelet transform set to 0.2, and number of colors set to 256. The first row represents methods without pre- or post-processing, the second row represents methods with clipping and SVD post-processing, and the third row represents methods with clipping and wavelet post-processing. The overall privacy guarantee for all perturbed FNC matrices is (*ϵ* = 2, *δ* = *N*^*—*3*/*2^)-DP. As expected, the Laplace mechanism performs worse than the analytic Gaussian mechanism since the *l*_1_ sensitivity is larger than the *l*_2_ sensitivity in high dimensions. Even with composition methods applied to reduce noise, Pure_k_ still does not outperform Approx. The Exponential mechanism is barely usable. Clipping and SVD post-processing significantly enhance the visual fidelity of private FNC matrices, whereas wavelet post-processing introduces additional visual distortion. This suggests that classical image processing techniques developed for natural images (e.g., photos) may not be suitable for this type of visualization.

This superior performance can be explained from both structural and theoretical perspectives. At a structural level, the privacy-preserving FNCs produced by this method retain the same block-structured organization seen in the original matrix, with recognizable within-network modules along the diagonal and comparatively weaker cross-network structure off-diagonal. The method preserves the relative ordering of strong versus weak connectivity patterns across domains, so the same major network-network relationships remain visually interpretable even though individual entries show slight deviations. In contrast, less effective DP examples introduce noise that partially blurs or fragments these modules, reducing contrast and making the within-network blocks less coherent, which can obscure higher-level structure.

From a theoretical standpoint, because the *l*_2_ sensitivity is lower than the *l*_1_ sensitivity in high dimensions, the analytic Gaussian mechanism adds less noise than the Laplace mechanism, resulting in better performance in terms of pattern preservation. The Exponential mechanism exhibits the worst visualization performance due to its high level of randomness. Specifically, the PDF of the output from Pure_k_ for each entry is proportional to 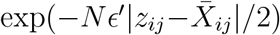, while the PDF of the output from Exp_k_ is proportional to exp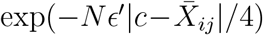. In general, the Exponential mechanism is more suitable for tasks such as determining product prices, where small perturbations in the determined price may lead to significant changes in revenue (utility). However, for FNC matrices, small changes in correlation do not substantially affect color assignment.

In terms of composition, exact composition and RDP-based composition outperform advanced composition, regardless of the mechanism used. For the Laplace mechanism, Pure_k_ requires less noise than Pure_r_, whereas for the analytic Gaussian mechanism, Approx_k_ requires more noise.

### 5.2. Processing Effect

We find that pre- and post-processing significantly enhance the visual utility of privacy-preserving data visualization. For FNC visualization, Figure 7g 7k and use different processing methods, and both exhibit the best visualization results, i.e., they are visually closest to the original average FNC shown in Figure 7a.

To quantitatively assess this observation, we first compute the MSE between the original average FNC and its private counterpart. As shown in Figure 8 (a)-(d), Figure A.1, and Figure A.2, the benefits of pre- and post-processing are more pronounced when the total privacy budget *ϵ* is small (high privacy) but become less significant as *ϵ* increases. The post-processing method based on SVD consistently achieves better performance than the wavelet transform.

While MSE captures pointwise numerical deviations, it does not reflect higher-level structural organization. Since FNC matrices are interpreted through block-structured functional networks, we additionally evaluate network modularity (Rubinov and Sporns, 2011). Modularity measures the strength of community structure relative to a random null model. A reduction in modularity after privacy perturbation indicates that intrinsic functional modules become less distinguishable, reflecting structural degradation. To further quantify local fluctuation patterns introduced by privacy noise, we compute the total variation (TV) (Michel et al., 2011) of the connectivity matrix and report the TV ratio between the private and original matrices. TV measures the aggregate magnitude of local differences between adjacent matrix entries, thereby characterizing high-frequency perturbations. A higher TV ratio indicates increased local irregularity and reduced visual smoothness.

As shown in Figure 8, Figure A.1, and Figure A.2, although SVD yields lower MSE across privacy budgets, wavelet-based post-processing better preserves structural properties and visual smoothness at larger *ϵ*. This divergence indicates that numerical, structural, and visual fidelity are not necessarily aligned.

**Figure 8.**
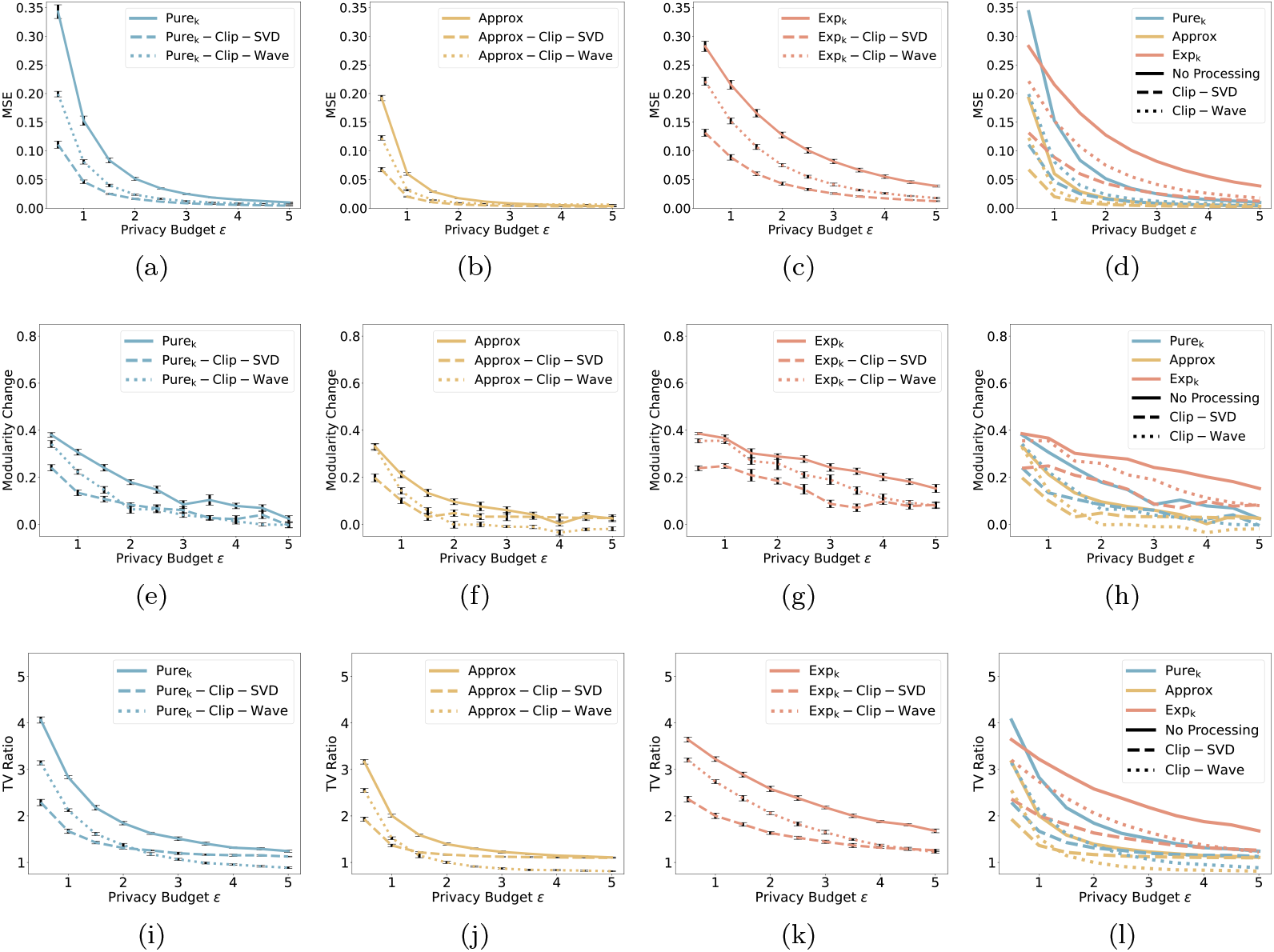
Quantitative evaluation of privacy-preserving FNC visualization on the FBIRN dataset. The first row reports MSE between the original and private average FNC matrices. The second row shows modularity change, defined as *Q*_orginal_ — *Q*_DP_, where *Q* denotes signed Louvain modularity. The third row presents the TV ratio, computed as TV_DP_*/*TV_orginal_. Columns correspond to (a)(e)(i) Pure_k_-, (b)(f)(j) Approx-, (c)(g)(k) Exp_k-_ related methods, and (d)(h)(l) all methods. Each configuration is repeated 50 times, and error bars are provided

For SBC visualization, pre-processing methods are not applicable. We apply the median filter technique as the post-processing method, resulting in a noticeable improvement in visualization quality (see Figure 6). These results suggest that different visualization types may require distinct pre- and/or post-processing methods.

### 5.3. Workflow Effectiveness

For connectogram visualization, we observe that our proposed workflow produces visualizations similar to its non-private counterpart (see Figure 9). Our work also finds that brain functional abnormalities in SZ are primarily located in the subcortical (SC), auditory (AU), sensorimotor (SM), visual (VI), and cerebellar (CB) domains Du et al. (2020). Given the importance of identifying these brain domains and regions rather than determining the exact connectivity values, we allocate more privacy budget to identify the brain domains and regions. Similar outcomes are observed in SBC visualization, where the privacy-preserving SBC map closely aligns with the SBC map without privacy protection. For instance, Figure 10 illustrates strong connectivities between the selected seed PCC and its neighboring voxels.

**Figure 9.**
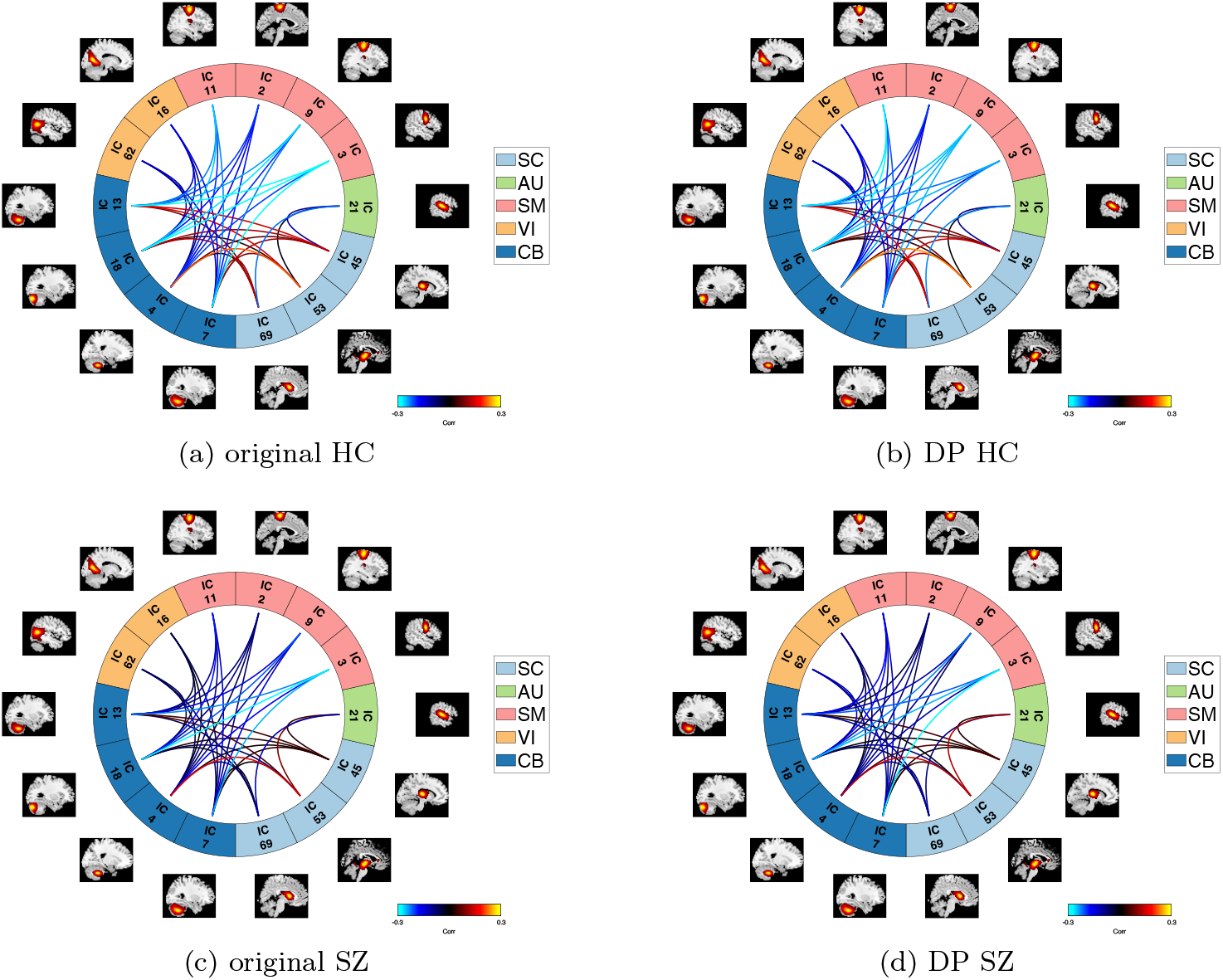
Privacy-preserving connectogram visualization with *ϵ* = 2, *p*_*d*_ = 5, and *p*_*r*_ = 14. We allocate 45% of the privacy budget to identify brain domains, another 45% to pinpoint brain regions, and reserve the remaining 10% to protect the connectivities.

**Figure 10.**
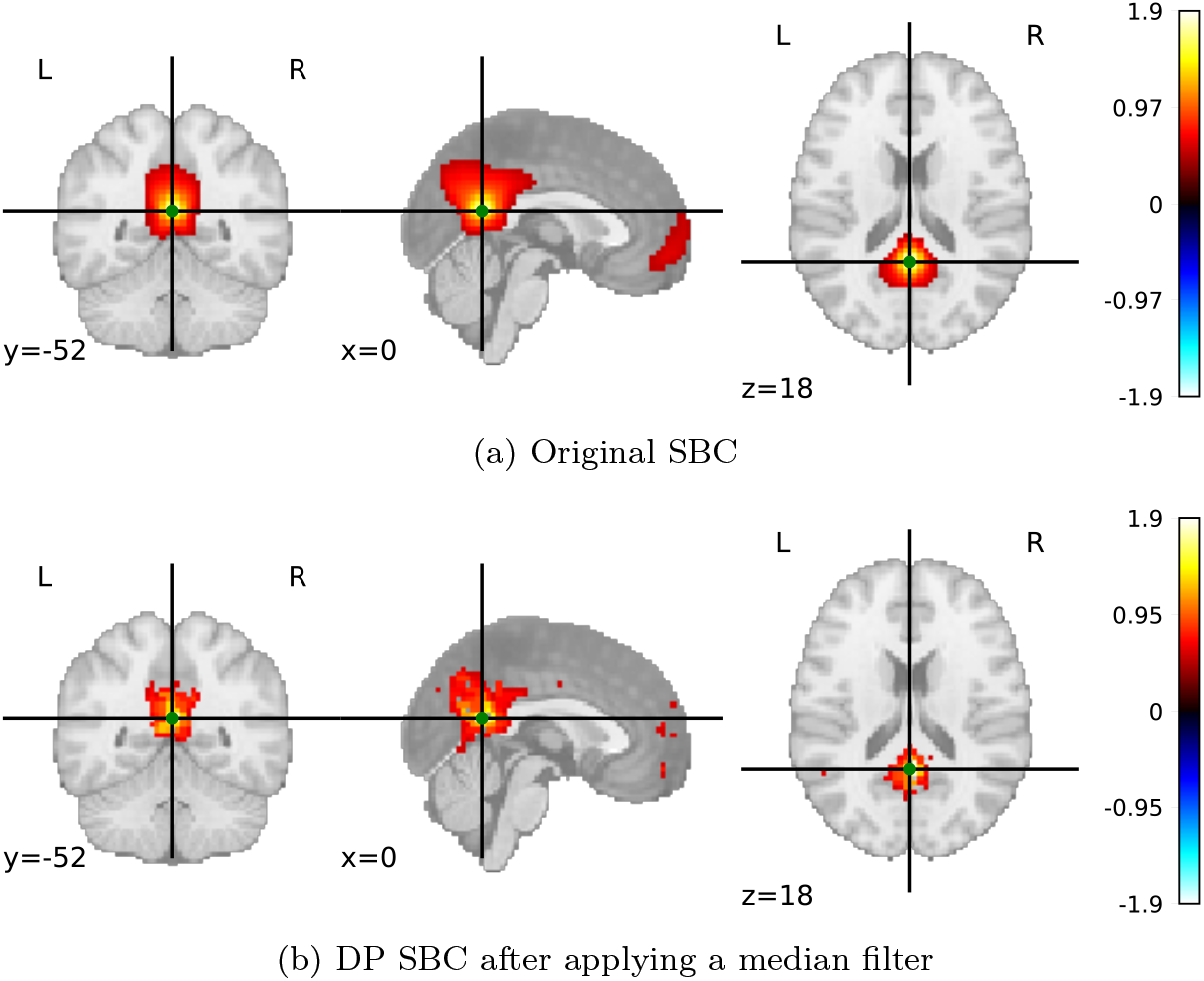
SBC maps on the NITRC dataset using the Posterior Cingulate Cortex (PCC) as the seed (indicated by the green dot), with parameters *N* = 1082, *ϵ* = 5, and *δ* = *N*^*—*3*/*2^. We allocate 50% of the privacy budget to identify significant correlations and 50% to protect the *z*-scores.

### 5.4. Visual Utility

It should be noted that there is no standard method for quantitatively assessing the quality of privacy-preserving visualizations. As discussed by Zhang et al. (2016), making a precise notion of “visual utility” is challenging. Errors are often measured using metrics such as the *l*_1_ or *l*_2_ norms, but these may not align with perceived quality. Moreover, in many cases, norm-based metrics may not be suitable for evaluating visualization quality. For example, in connectogram visualization, hypothesis testing errors (seeing irrelevant domains or regions, or failing to notice significant ones) may provide a more relevant evaluation approach, as the primary concern is identifying domains or regions that exhibit significant differences in group comparisons. When it comes to SBC visualization, identifying quantitative methods to assess visual utility becomes even more challenging.

We surveyed neuroimaging researchers (at various levels of experience) through a convenience sample of researchers interested in functional network connectivity. We designed a survey consisting of five sections and collected 22 responses, with 18 participants consenting to the release of their responses. The full questionnaire, participant responses, and a summary of survey results are provided in the supplementary material. Our findings from this initial survey are as follows:

- Algorithm comparison: Participants found Approx-based methods, especially those with clipping and SVD, most similar to the original. In contrast, wavelet-based versions, although achieving lower MSE, were frequently rated as more visually distorted. This suggests that quantitative metrics like MSE alone may not fully capture perceptual quality.
- Privacy–utility tradeoff: As the privacy budget *ϵ* increased, visual quality tended to improve, showing clearer structures and fewer distortions.
- Consistency under the same privacy level: Most participants felt that visualizations generated using the same algorithm and privacy level were generally similar, despite differences in noise realizations.
- Connectogram workflow evaluation: Participants observed that allocating more privacy budget to brain region (node) identification improved perceived visualization quality. Conversely, reducing the budget for nodes increased visual distortions. This effect was not primarily caused by node misidentification itself but rather because node errors propagated to edges, resulting in larger and more noticeable changes in the central visual area.
- SBC workflow evaluation: Participants generally considered the privacy-preserving visualization usable and effective at preserving core connectivity patterns. However, some noted reduced smoothness and missing functional connections, suggesting that a higher privacy budget may be needed to better retain critical connections.

We believe this initial feedback provides some guidance for future research directions. In particular, we believe that for privacy-preserving visualization to become part of the regular workflow in future software systems for neuroimaging research, it is important to assess the visual utility as judged by the intended audience for the visualization.

## 6. Future Directions

Data visualizations are often designed to provide qualitative assessments of quantitative data. We discussed the risks of sensitive information leakage in visualizations, highlighting the growing importance of privacy protection in this area. Current research has identified various privacy attacks across fields such as machine learning (Rigaki and Garcia, 2023). Adversaries may seek to access sensitive information, including the training dataset, model parameters, hyperparameters, and architecture, etc. Inspired by these concerns, we believe that privacy attacks in visualizations are also an area worth exploring.

When visualizing sensitive data, the gap between qualitative assessments and quantitative precision offers many interesting options for guaranteeing privacy. In this study, we explored several approaches to providing privacy for visualizations related to functional network connectivity. We view this as a *starting point* for considering privacy in data visualization within neuroimaging and human health research more broadly. In these settings, individual data holders may not have a large sample size but in systems using federated learning, the differences among these methods would be much smaller as they would require less noise and provide better utility due to the increased sample size.

We did not explore the full gamut of postprocessing methods or transform-based privacy algorithms. For example, alternative processing methods such as Fourier transforms and bit plane slicing could be explored. Another approach involves transforming data to obtain sparse coefficients, applying privacy-preserving techniques to protect them, and then using the inverse transform to reconstruct the privacy-preserving data. In addition to refining these approaches, manipulating the choice of color map, the number of colors | 𝒩|, and the quantization scheme can all potentially improve the visual effect. Many visualizations seek to represent the uncertainty in estimates or observations. Uncertainty visualization has long been recognized as a critical component for scientific communication, with numerous studies highlighting its importance (MacEachren, 1992; MacEachren et al., 2005; Potter et al., 2010). Visualizing uncertainty and providing privacy can potentially present synergies: designing effective algorithms is an interesting avenue for future work. A starting point would be to develop privacy-preserving displays for error bars and confidence intervals. When privacy is included, we might find that other visual representations of uncertainty are more visually salient.

## Data and Code Availability

This study utilizes publicly available datasets accessible directly from their respective repositories; signing a Data Use Agreement (DUA) may be required. The code is available from the corresponding author upon reasonable request.

## Conflicts of Interest

The authors declare no conflicts of interest that may bias or could be perceived to bias this work.

## Acknowledgments

The work of authors was supported in part by the US National Institutes of Health under awards 2R01DA040487, R01DA049238, and R01MH121246, as well as by the National Science Foundation under award 2112455. The work of Sarwate was also supported by the National Science Foundation under award CNS-2148104 and is supported in part by funds from federal agency and industry partners as specified in the Resilient & Intelligent NextG Systems (RINGS) program.

The authors wish to thank the FBIRN, BLSA, and ADNI teams. In addition, the authors acknowledge the Neuro Bureau, the ADHD 200 consortium, and Virginia Tech’s ARC. The authors also acknowledge the assistance of ChatGPT for English language editing.

## Appendix

The appendix contains additional details and figures omitted from the main manuscript.

**Figure A.1.**
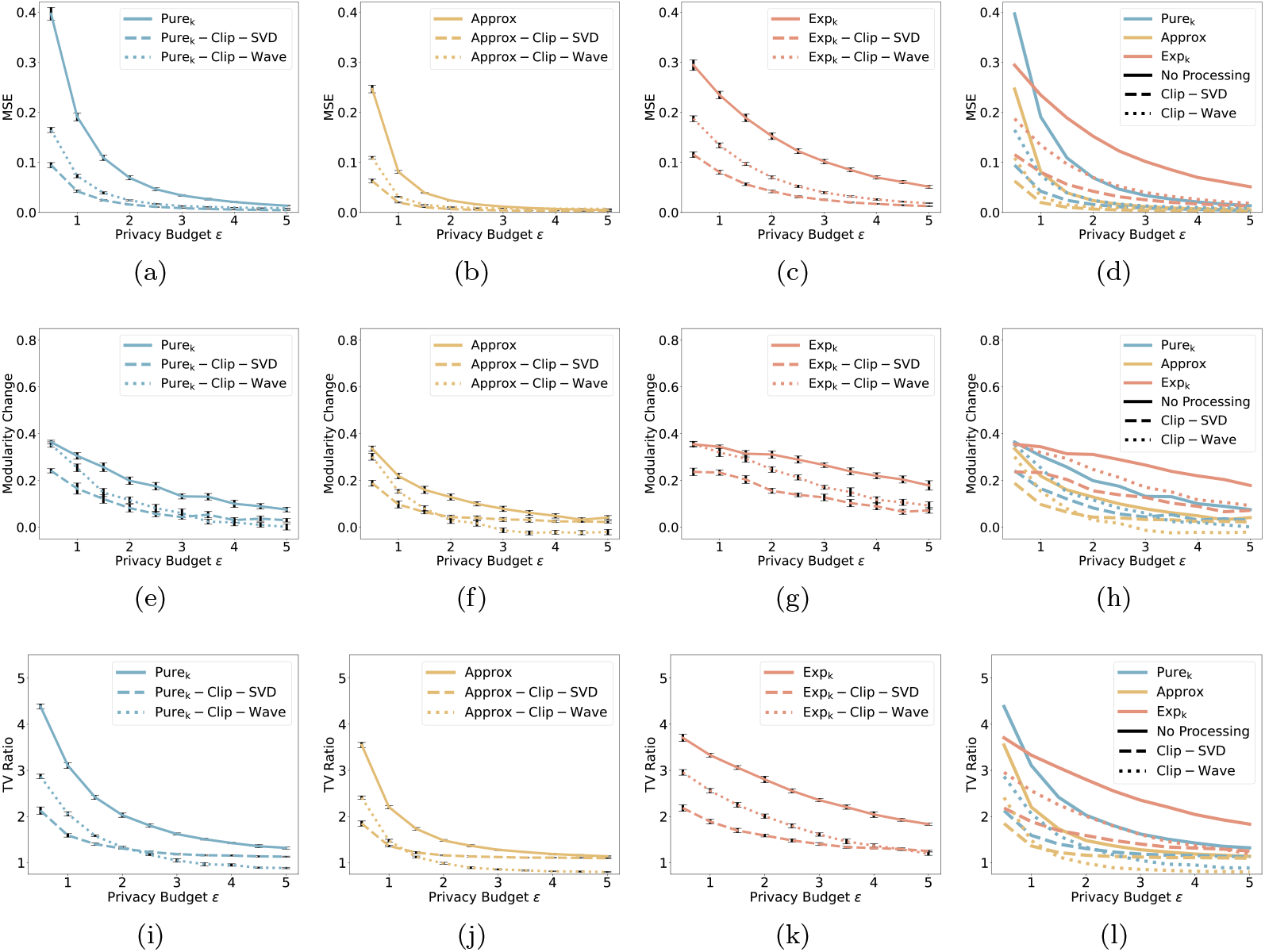
Quantitative evaluation of privacy-preserving FNC visualization on the BLSA dataset. The first row reports MSE between the original and private average FNC matrices. The second row shows modularity change, defined as *Q*_orginal_ — *Q*_DP_, where *Q* denotes signed Louvain modularity. The third row presents the TV ratio, computed as TV_DP_*/*TV_orginal_. Columns correspond to (a)(e)(i) Pure_k_-, (b)(f)(j) Approx-, (c)(g)(k) Exp_k-_ related methods, and (d)(h)(l) all methods. Each configuration is repeated 50 times, and error bars are provided.

**Figure A.2.**
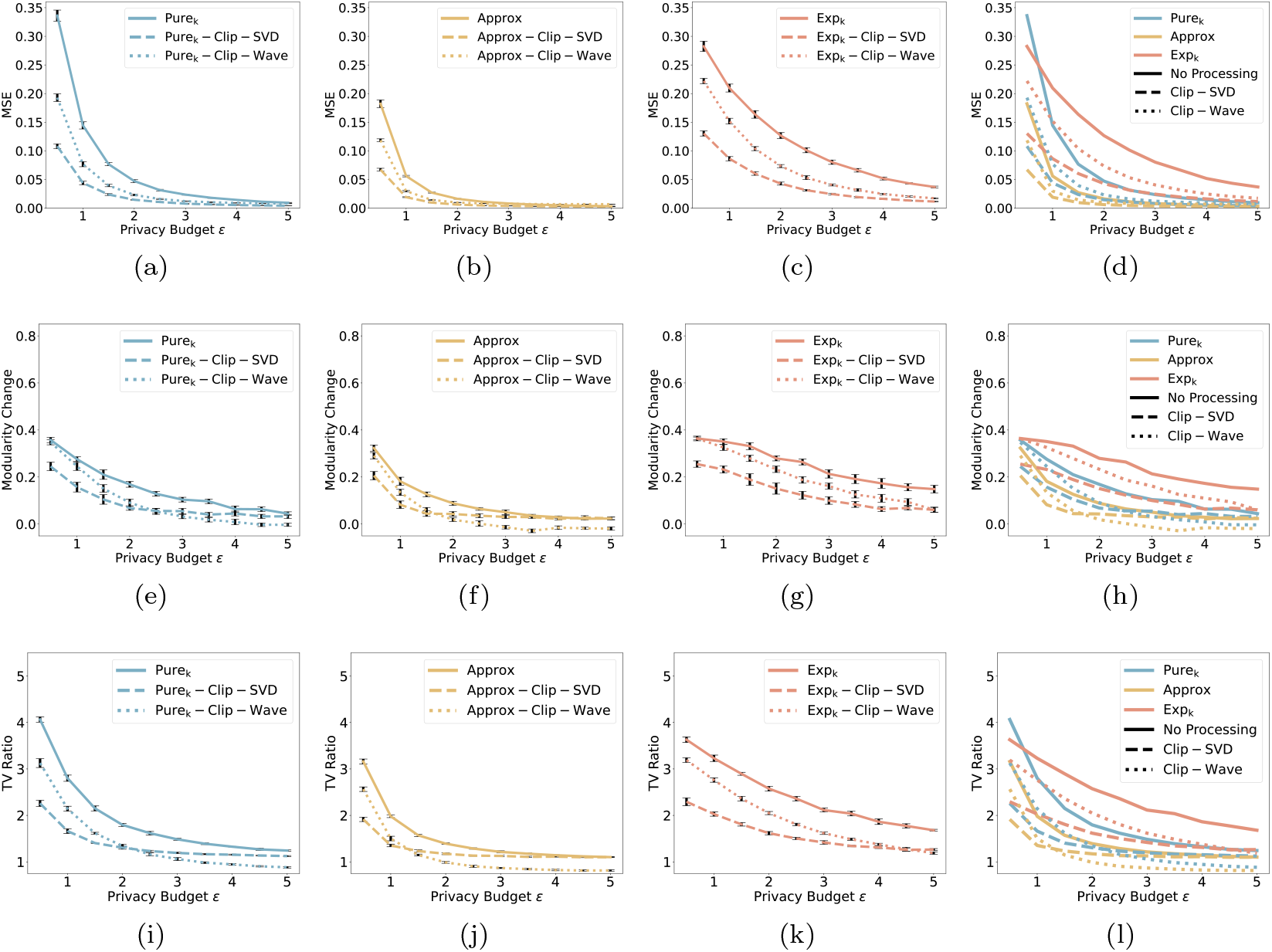
Quantitative evaluation of privacy-preserving FNC visualization on the ADNI dataset. The first row reports MSE between the original and private average FNC matrices. The second row shows modularity change, defined as *Q*_orginal_ — *Q*_DP_, where *Q* denotes signed Louvain modularity. The third row presents the TV ratio, computed as TV_DP_*/*TV_orginal_. Columns correspond to (a)(e)(i) Pure_k_-, (b)(f)(j) Approx-, (c)(g)(k) Exp_k-_ related methods, and (d)(h)(l) all methods. Each configuration is repeated 50 times, and error bars are provided.

**Figure A.3.**
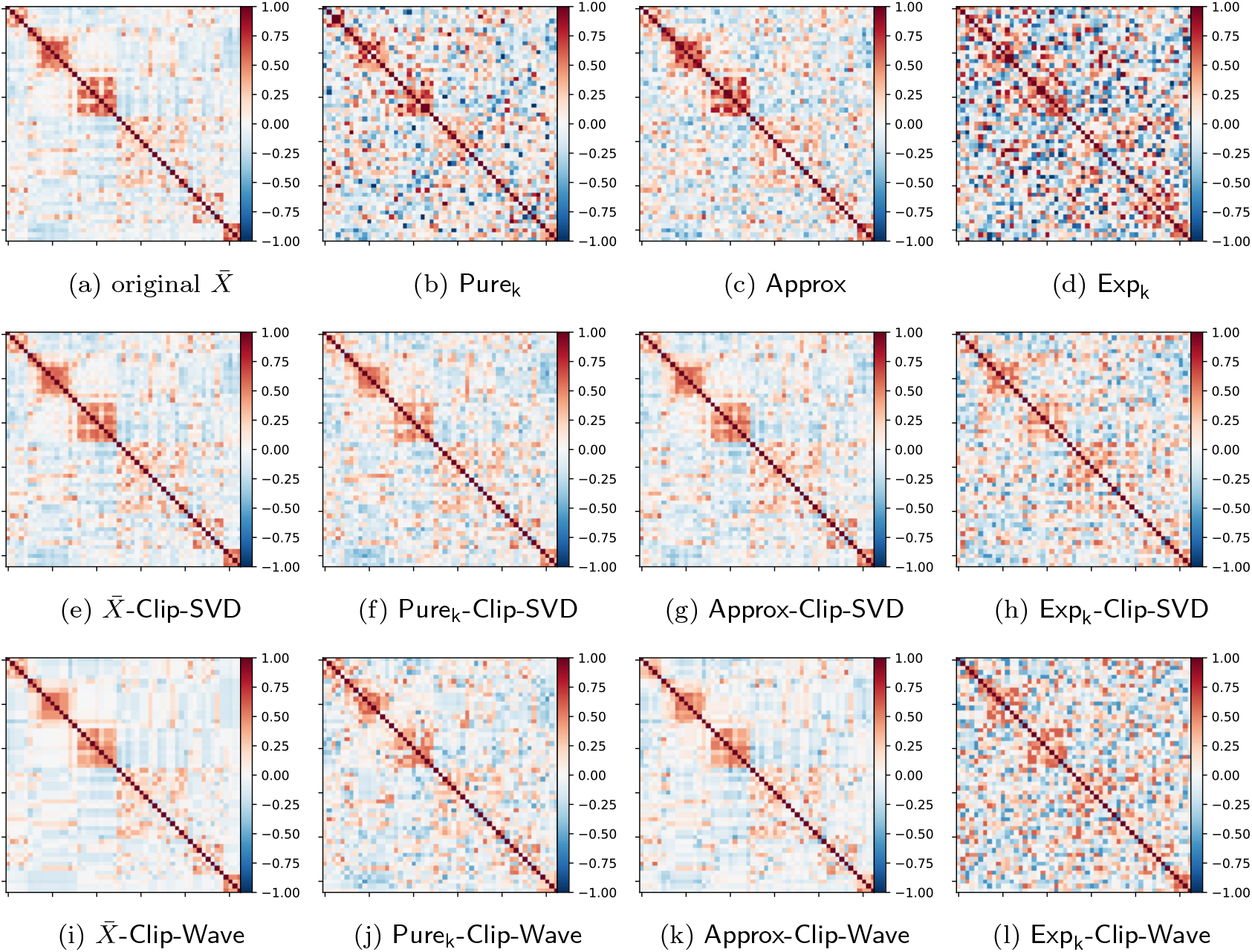
Visualization of the BLSA dataset with *N* = 891, *ϵ* = 2, *δ* = *N*^*—*3*/*2^, clipping bound *b* = 0.7, number of components used in SVD set to 10, threshold used in the wavelet transform set to 0.2, and number of colors set to 256.

**Figure A.4.**
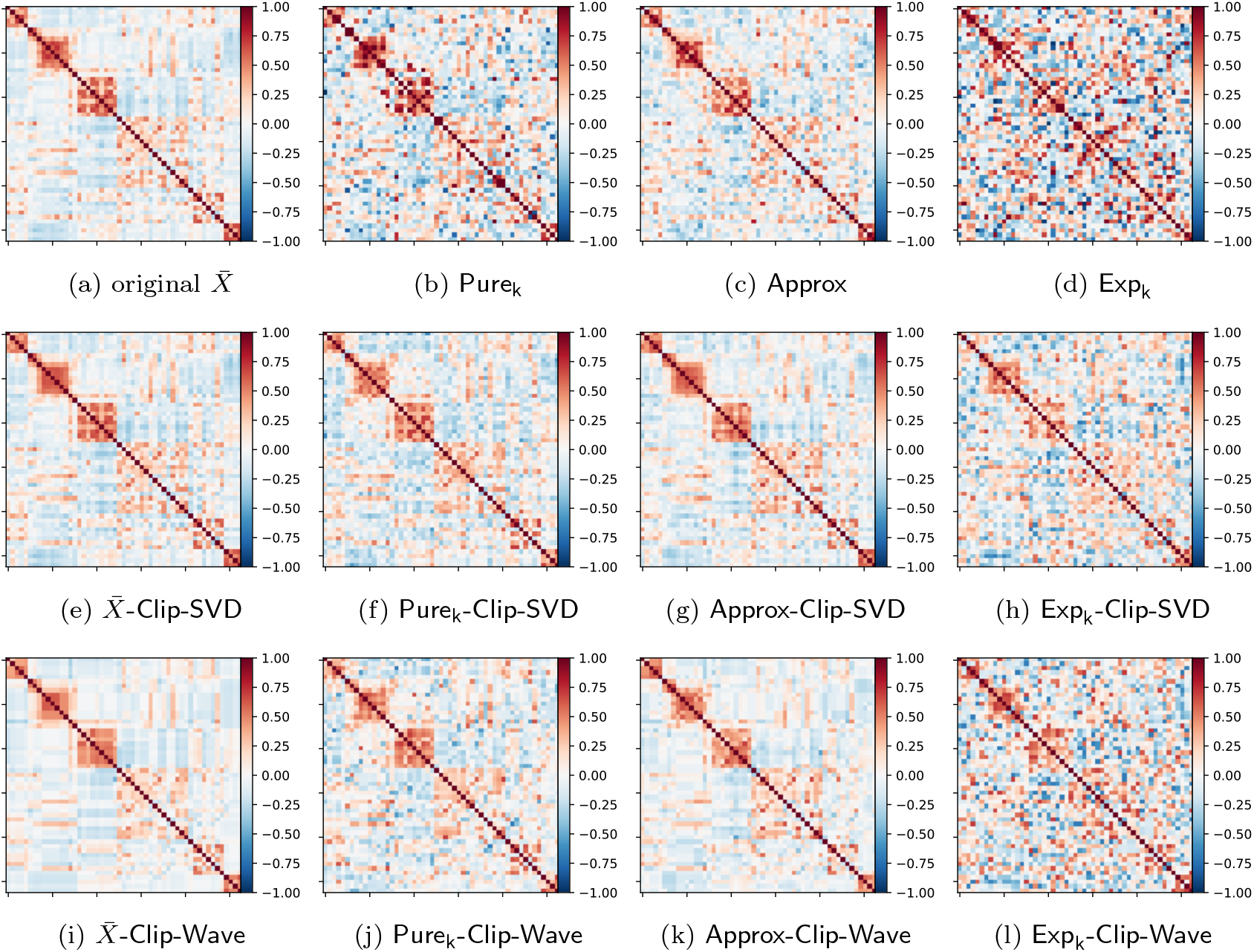
Visualization of the ADNI dataset with *N* = 1091, *ϵ* = 2, *δ* = *N*^*—*3*/*2^, clipping bound *b* = 0.75, number of components used in SVD set to 10, threshold used in the wavelet transform set to 0.2, and number of colors set to 256.

## Notes

### Competing Interest Statement

The authors have declared no competing interest.

### Summary of Updates

The manuscript has been revised to improve clarity and presentation. The Introduction has been revised to better clarify the motivation of the study. Additional quantitative evaluation metrics beyond MSE have been included, and an expert assessment has been added to complement the evaluation. Figures have also been updated to vector format.

